# Toward welfare indicators: leg condition and aversive stimuli modulate defensive behavior in Opilones

**DOI:** 10.64898/2026.10.29.762782

**Authors:** Shannon L. O’Brien, Marcelo Araya-Salas, Noah Skelly, Dulce M. Herrera Palencia, Ignacio Escalante

## Abstract

The history of invertebrate use in research and their growing use in managed settings highlight the need for validated behavioral indicators of welfare, yet these remain poorly understood across taxa. The arachnid group Opiliones (“harvestmen” or “daddy long-legs”) is an ideal group for identifying welfare indicators, given their well-characterized defensive behavior, including the common strategy of leg autotomy. The goal of this project was to identify candidate behavioral welfare indicators for *Prionostemma* sp. 1 in the Neotropical rainforest. We brought field-collected individuals with different leg conditions to the lab and briefly exposed them to an aversive stimulus (a gentle squeeze and release) paired with an assigned substrate (either rock or sand). A rewarding stimulus (food) was provided on the alternate substrate. We found no evidence of preference for roosting on the substrate associated with the rewarding stimulus, nor avoidance of the substrate associated with the aversive stimulus. Thus, evidence for associative learning was not supported. Additionally, substrate use did not differ based on leg condition (intact, missing 1-2 legs, or missing sensory legs). After an aversive stimulus, fleeing was consistently the most common response throughout 5 days. In contrast, the frequency of bobbing increased while the frequency of escaping decreased over trials. Individuals with all legs tended to escape more often than those missing legs, but neither the number nor the type of missing legs substantially improved predictions of behavioral responses. Overall, fleeing and bobbing may serve as behavioral indicators of negative welfare states, whereas the observed absence of exploratory tapping following the aversive stimulus suggests that this behavior may indicate neutral or positive welfare states. As one of the first investigations of welfare indicators in wild arachnids, this study provides novel insights into how defensive behaviors and physical condition may inform assessments of invertebrate welfare, laying the groundwork for future validation of behavioral indicators.

## Introduction

Invertebrates comprise most animal biodiversity and are increasingly utilized within managed contexts, such as zoos and aquariums, research laboratories, classrooms, as well as within the micro-livestock and aquaculture industries (O’Brien et al. 2026; Barrett and Adcock 2023; Wilson-Sanders 2011). This growing use has broadened interest in their welfare (Crump et al. 2023; Lewbart and Zachariah 2023; Wahltinez et al. 2022; Keller 2017), driving a notable rise in welfare-focused invertebrate research in recent years (e.g., Fossati et al. 2024, Rowe et al. 2024; Crook 2021), especially for cephalopod mollusks and decapod crustaceans in both managed and wild contexts (Browning et al. 2025; Andrade et al. 2023, Albalat et al. 2022, Cooke et al. 2019). However, research on invertebrate welfare remains outpaced by welfare research focused on vertebrate taxa (Binding et al. 2020), a pattern that mirrors research in animal behavior (Rosenthal et al. 2017) and evolution (Bacon et al. 2023). Emerging evidence indicates that at least some invertebrate taxa are likely sentient and capable of experiencing both negative and positive affective states (Romero-González et al. 2025; Crump et al. 2022; Galpayage Dona et al. 2022). Notable advancements stemming from this work include the inclusion of cephalopod mollusks and decapod crustaceans within the UK’s Animal Welfare Sentience Act of 2022 (Animal Welfare (Sentience) Act 2022) and the inclusion of these taxa as well as insects within the New York Declaration on Animal Consciousness (Andrews et al. 2024). Recent work proposes that even without definitive evidence of sentience, precautionary approaches and welfare-centered practices should be prioritized (de Souza Valente 2025, Fischer et al. 2025).

Efforts to evaluate invertebrate welfare face considerable methodological challenges. Established welfare frameworks, such as the Five Freedoms (FAWC 1993) and the Five Domains (Mellor et al. 2020), have been developed with vertebrates in mind and rely on well-characterized links between physiology, behavior, and affective states in those taxa (Barrett and Fischer 2023). In contrast, invertebrates have highly diverse nervous systems, sensory modalities, and behavioral repertoires, making it difficult to infer affective states using existing frameworks (Barrett and Fischer 2023; Perkins 2021). One welfare assessment approach, the Animal Welfare Assessment Grid (AWAG, Justice et al. 2017), has been adapted to assess welfare in cockroaches (Free and Wolfensohn 2023) as well as decapods and cephalopods (Narshi et al. 2022) in managed care. Additionally, a welfare assessment tool geared towards invertebrates used as feed in zoos and aquariums has been developed (O’Brien et al. 2026), highlighting recent advancements in this field. Advances such as these allow us to make evidence-based decisions that support the welfare of invertebrates in human care. However, further research is needed to develop direct animal-based indicators of welfare (e.g., physiological or behavioral measures) for most invertebrate taxa. Similarly, research on the welfare of wild invertebrates is in its early stages and warrants further investigation. Studying invertebrates in their natural environments can provide insight into the ecological and evolutionary factors shaping behavioral responses to challenges and opportunities, and how these responses may relate to welfare. Such research can also help establish species-typical behavioral baselines, identify candidate indicators of welfare, and clarify how environmental conditions and individual characteristics influence welfare outcomes. These insights can inform welfare assessments in managed settings while advancing our understanding of the factors influencing the welfare of wild invertebrate populations.

One promising approach to advancing invertebrate welfare science is to focus on studying organisms whose natural history and behavior are well described. Some species of the arachnid order Opiliones (commonly known as “harvestmen”, “harvesters”, or “daddy long-legs”) represent one such system (e.g., Machado and Burns 2024; Escalante and O’Brien 2024; Giribet and Sharma 2015; Willemart et al. 2009). Opiliones, particularly the ones in the suborder Eupnoi, are characterized by their elongated legs and a reliance on multiple defensive strategies, including chemical and mechanical defenses (Escalante and O’Brien 2024; Freire et al. 2026; Raspotnig et al. 2022; Powell et al. 2021; Pinto-da-Rocha et al. 2007; Ximenes and Willemart 2024). One notable defense is their ability to voluntarily release legs, known as autotomy (Guffey 1999; Emberts et al. 2019; Kinoshita et al. 2026; Powell et al. 2021). In natural populations, approximately half of the individuals are missing one or more legs (Escalante and O’Brien 2024). Previous studies have demonstrated that Opiliones exhibit robustness to leg loss, with relatively limited or short-term impacts on locomotion, physiology, mating success, and survival (Escalante et al. 2020; 2021; Escalante and Elias 2021; 2022). However, the impact of autotomy on welfare remains unexplored. Understanding these effects will be valuable for developing a more comprehensive view of how invertebrates cope with changes in bodily condition, such as leg loss, which often arise from environmental challenges such as predation pressure and even disease (Maginnis 2006; Emberts et al. 2019). Thus, Opiliones are ideal organisms for exploring the welfare implications of autotomy and deepening our understanding of invertebrate welfare more broadly.

The first aim of this project was to quantify behavioral responses to a brief exposure to an aversive stimulus in individuals of one species of Opiliones (*Prionostemma* sp. 1; Fig. 1A, B). By documenting the frequency and type of responses expressed under these conditions, we aimed to identify candidate behavioral indicators of negative welfare states. We predicted that individuals would exhibit increased defensive behaviors, such as fleeing, escaping, and bobbing, following exposure to an aversive stimulus. We further predicted that behavioral responses would differ according to leg condition, as individuals missing legs may experience altered vulnerability or physical constraints that influence their defensive strategies. Specifically, we expected individuals missing legs to exhibit different frequencies or types of defensive responses compared to individuals with all legs intact. Conversely, species-typical behaviors that are reduced or absent following exposure to an aversive stimulus may represent candidate indicators of neutral or positive welfare states.

**Figure 1.**
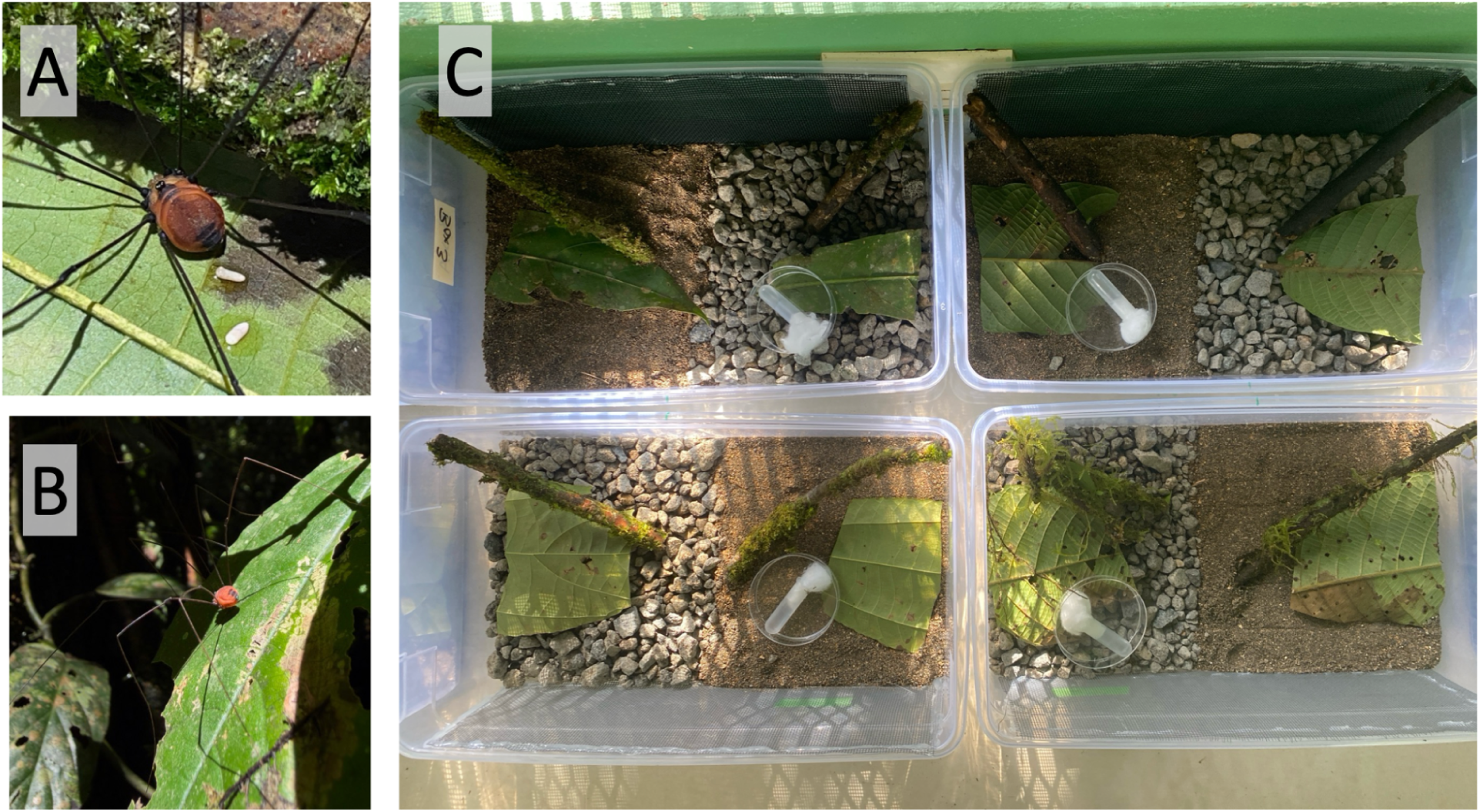
Study organisms and experimental settings. A. close up of *Prionostemma* sp.1 Opiliones. B. One of the study animals in their natural habitat before being collected. C. Housing conditions for the experiments. Lidded plastic containers (12x7x4 inches) were filled half with sand and half with small rocks. La Selva Research Station, Sarapiquí, Costa Rica, 2026.

The second aim was to assess evidence for associative learning, which could provide important insights into how individuals perceive and respond to their environment. Previous work has demonstrated learning abilities in Opiliones (Pagoti et al. 2025; dos Santos et al. 2013), suggesting that these animals can modify their behavior based on experience. Building on this foundation, we conducted paired substrate-stimulus trials to test whether individuals, including those with all legs intact and those missing different numbers and types of legs—Opiliones have two secondarily modified sensory legs—could learn associations between distinct substrates and either an aversive or a rewarding stimulus. We predicted that individuals would demonstrate associative learning by preferentially selecting the substrate associated with the rewarding stimulus and/or avoiding the substrate associated with the aversive stimulus. We also explored whether leg condition influenced associative learning, as individuals missing legs may experience different costs and benefits associated with environmental risks and rewards. If leg loss increases perceived vulnerability, we predicted that individuals missing legs would demonstrate stronger avoidance of the substrate associated with the aversive stimulus compared to individuals with all legs intact.

Demonstrating learning as predicted would strengthen evidence that Opiliones use environmental cues to guide decision-making, a capacity with important implications for both ecology and animal welfare. In natural environments, risks and benefits may shift for individuals missing legs. The ability to adjust behavior based on learned associations could facilitate adaptive responses to changes in bodily condition, such as modifying habitat use to mitigate presumed vulnerability (e.g., Escalante and Elias 2021).More broadly, understanding how Opiliones respond to stimuli, how these responses vary with leg condition, and their capacity for associative learning can inform the development of behavioral welfare indicators and advance evidence-based approaches to invertebrate welfare research, management, and policy in both wild and managed settings.

## Methods

### Study Species

We conducted this research with adult Opiliones of *Prionostemma* sp.1 (family: Sclerosomatidae) (following the nomenclature in Proud et al. 2012 and Freire et al. 2026), which we collected from the Tres Ríos Trail (STR) at La Selva Research Station in Puerto Viejo de Sarapiqui, Costa Rica (10° 26′ N, 84° 00′ W, 50 m elevation). We looked for individuals during the morning (08:00 to 11:00 hours), when they roost on tree trunks, buttresses, and leaves of palms and Heliconiales (Escalante et al. 2019, Villaseñor-Amador & Escalante 2025, 2026).

We conducted this work with 50 intact (all 8 legs present) individuals we found in January 2026 and 9 intact and 48 autotomized individuals in June 2026, for a total of 107 individuals. Autotomized individuals were those missing one (n=29) or two (n=19) legs in total. The subset of intact individuals tested in June 2026 enabled us to control for potential temporal differences in the responses of these Opiliones (see below for further details).

### Animal Housing and Husbandry

Animals were housed individually in lidded plastic containers (12 x 7 x 4 inches) in an ambient, open-air field lab that provided natural light, temperature (25.8 ± 1.66 °C), and relative humidity (90.9 ± 3.3 %). The bottom of each container was filled with sand on one side and small rocks on the other side (Fig. 1C). One fresh leaf and one mossy twig were placed on each side of the container to provide substrate roughness for climbing and adequate humidity. Fine mesh was adhered to one side of the container to provide additional climbing surface; the mesh extended across the long side of the container, so it was available on both the sand and rock sides. The center of the plastic lid was removed and replaced with a fine mesh to provide airflow and an additional climbing surface.

Animals were provided with *ad libitum* quantities of fresh fruit and a mixture of commercial pet foods (e.g., crested gecko pellets, bearded dragon pellets, and fish flakes), as done in previous research (Sasson et al. 2020; Brown et al. 2025; Schoffstall et al. 2013). Water was provided in a test tube topped with a cotton ball for slow dispensation (Acosta & Machado 2007; Sasson et al. 2020). Food and water were replaced daily at 19:00 hours to ensure freshness. Leaves were replaced as needed when they became dry (approximately every 2-3 days).

### Lab Experiments

Animals were allowed to acclimate for 3 full days following collection. During this time, animals were not disturbed except once daily, in the evening, to replace their food and water. During the acclimation phase, each animal’s substrate use was recorded three times daily (8:00, 14:00, 19:00) to assess whether individuals had any inherent substrate preferences prior to training.

Following acclimation, animals were trained to associate one substrate with a rewarding stimulus and the other substrate with an aversive stimulus (substrates systematically alternated: half of the individuals were trained to associate rock with the aversive stimulus and sand with the rewarding stimulus, the other half of the individuals were trained to associate sand with the aversive stimulus and rock with the rewarding stimulus). During the training phase, animals were trained twice daily (8:00 and 14:00) for five days. Previous research with Opiliones suggests they can learn within this timeframe (dos Santos et al. 2013). During training, we first recorded the substrate on which the animal was located (as during the acclimation phase). Then, each individual was collected from their home container by their back two legs by researcher 1 (I.E.) and gently squeezed on their abdomen between forefinger and thumb (aversive stimulus). Simultaneously, researcher 2 (S.L.O., N.S., or D.M.H.P.) placed fresh food pellets on the substrate associated with food (rewarding stimulus). The animal was then released facing forward, dorsal side up, on the substrate associated with the aversive stimulus.

Each animal’s behavioral response during the 5 seconds after release was recorded. This timeframe was chosen after pilot trials showed that the animals settled and returned to a neutral posture within 5 seconds after release. The five behaviors of interest recorded were: bobbing, freeze, flee, escape, and chemical release (see full description on the ethogram in Table 1, and Supplementary Movie 1). The experimenter handling the animals cleaned their gloves after each trial with a cotton ball soaked in ethanol and changed the gloves every 5 trials, or immediately after a trial if the animal released the defensive chemical secretion (as done in Freire et al. 2026).

**Table 1.** Behavioral ethogram for Opiliones of *Prionostemma* sp.1 (Sclerosomatidae).

| Behavior | Description |
| --- | --- |
| Bobbing | The animal repeatedly moves its body up-and-down by extending and contracting multiple legs. Bobbing can occur when the animal is moving across the terrarium or while remaining in the same location (Holmberg et al. 1984, Escalante et al. 2019, Villaseñor-Amador and Escalante 2025, Calhoun et al. 2025). |
| Freeze | The animal becomes immobile and remains stationary in an upright posture (distinct from “thanatosis” described in Powell et al. 2021). |
| Flee | The animal moves away from the point of release by running and/or walking (Holmberg et al. 1984, Escalante et al. 2019) but remains within the confines of their home container. |
| Escape | The animal moves and runs away (Escalante et al. 2019) from the point of release, exiting the confines of their home container. Note: when this occurred, data collection stopped, and the animal was immediately returned to their home container. |
| Chemical release | The animal releases a volatile milky secretion from the dorsal openings (ozopores) of their repugnatory glands (Escalante 2019, Ximenes & Willemart 2024, Freire et al. 2026). The secretion was visually confirmed by observing the liquid on their dorsum and by smelling it (following the same procedure as in Freire et al., 2026). |
| Autotomy (leg loss) | The animal releases one (or more) leg(s) quickly. The leg detaches from the coxae-throcanther joint and twitches for >10 seconds (reviewed in Escalante & O’Brien 2024). |
| Leg tapping | Constant and fast up-and-down and sideways movements of the distal half of the legs of Opiliones (Fowler-Finn et al. 2014, Dias et al. 2014). Tapping is more commonly performed by the legs of the second pair, the secondarily modified antenniform leg pair, followed in frequency by the legs of the first pair. |
| Grooming | The animal bends a given leg and gets it in proximity of the mouthparts (chelicerae and pedipalps). The mouth parts grab the leg by the distal half. The animal then cleans the leg by gently pressing its mouthparts against it and moving the leg away from the mouth (Pereira et al. 2004, Willemart et al. 2006). The leg forms an irregular loop that grows in diameter as the animal approaches the tip of the leg to the mouth. |
| Feeding | Upon touching the food source (in this case, the small Petri dish with food items), the animal stops and lowers its body so its mouthparts contact the food. The animal’s proximal side remains close to the food while the pedipalps cut and grab small chunks of food, usually spinning the item. Then the pedipalps carry the items to the chelicerae, which move up and down before ingestion. |

During the five training days, each animal’s substrate use was recorded three times daily (8:00, 14:00, 19:00) to assess whether individuals’ substrate use changed as a result of training (e.g., the animal recorded more often on the substrate associated with a rewarding stimulus or recorded less often on the substrate associated with the aversive stimulus). Substrate use was recorded prior to disturbing the animals for training and feeding.

### Data Analyses

Substrate use and behavioral responses were analyzed using Bayesian generalized linear mixed models fitted in the R package *brms* (v2.23.0; Bürkner 2017; 2018), which interfaces with Stan (Stan Development Team 2021) to perform Hamiltonian Monte Carlo (HMC) sampling from the posterior distributions of model parameters. Each observation corresponded to a unique combination of individual and sampling period (each observation or trial bout across days). Because all 107 individuals were observed repeatedly across multiple sampling periods, individual identity was included as a random intercept in all models to account for non-independence among observations from the same individual. Field season (January or June 2026) was included as a fixed effect in all models to account for potential differences across seasons. Sampling period and number of legs (8, 7, or 6) were modeled as monotonic effects, which allow ordered predictors to have potentially unequal changes between consecutive levels while retaining their ordinal structure (Bürkner & Charpentier 2020).

For each response, we used a targeted candidate-model approach to evaluate the predictive contribution of the biological predictors, rather than exhaustively comparing all possible combinations of predictors. Candidate models were compared using approximate leave-one-out cross-validation (LOO; Vehtari et al. 2017; 2024), implemented in the R package *loo* (v2.10.1; Vehtari et al. 2025). LOO is a predictive model-comparison approach analogous in purpose to information criteria such as Akaike’s (AIC), but evaluates models based on their expected out-of-sample predictive performance while accounting for uncertainty in the fitted models. Models were ranked by their expected log predictive density (elpd), with the model having the highest elpd considered to have the greatest predictive support. We used differences between models (delta elpd) to assess the relative predictive contribution of individual predictors and to evaluate the biologically motivated interaction hypotheses described below.

To identify which levels of the categorical predictors showed differences, we calculated posterior inferences based on model-derived predicted probabilities and posterior contrasts, using the R package *marginaleffects* (v0.32.0; Arel-Bundock et al. 2024). Predicted probabilities represent the estimated probability of each response category under specified predictor values. Posterior contrasts quantify the expected change in predicted probability associated with changing one predictor while averaging over the observed values of the remaining predictors. These contrasts are analogous to estimated marginal mean contrasts commonly used in frequentist analyses but are derived directly from the posterior distribution.

All posterior estimates are reported as posterior means with 95% uncertainty intervals. All models were fitted using the default weakly informative priors implemented in *brms*, intended to regularize estimation while remaining weakly informative with respect to the expected magnitude of biological effects. Models were run for 4,000 iterations per chain, with the first 2,000 iterations discarded as a warm-up. Model convergence was assessed using the potential scale reduction factor (R-hat), effective sample sizes, and number of divergent transitions. Posterior distributions were additionally inspected visually using trace plots.

### Substrate Use Modeling

Substrate use was analyzed using a Bayesian categorical generalized linear mixed model. The response variable was preferred substrate, with three categories: rock, sand, and other. For rock and sand, we included observations of an individual roosting directly on the substrate, as well as on twigs or leaves located on that substrate. The “other substrate” category included observations on the mesh or at the very center of the rock-and-sand area. Candidate fixed effects included sampling period, food substrate (i.e., the substrate on which food was presented), number of legs (8, 7, or 6), sensory leg condition (both sensory legs present or missing one or two sensory legs), and experimental phase (acclimation or training phase). We also evaluated two biologically motivated interaction hypotheses. A first interaction between the experimental phase and substrate tested whether any observed changes in substrate use corresponded with training (such that individuals spent more time on the substrate associated with the rewarding stimulus or less time on the substrate associated with the aversive stimulus). This interaction would provide evidence for associative learning in Opiliones. A second interaction between the experimental phase, substrate, and leg condition (three-way interaction) tested for differences in substrate preference across time between individuals with all legs intact and those missing one or more legs.

In total, 17 competing models were fitted: one full additive model containing all biological predictors, one sampling-period-only model, four single-predictor models, four sampling-period-plus-single-predictor models, one full model without sampling period, four leave-one-predictor-out models, and the two interaction models described above. The full additive model was among the best-supported models (see Results and Supplementary Table 1); neither interaction improved predictive performance relative to the full model. Hence, we used the full additive model for biological inference. For categorical predictors, posterior contrasts were used to estimate differences in the probability of using each substrate between predictor levels (Supplementary Table 2). Because the sampling period was modeled as a monotonic effect, temporal changes in substrate use were instead described using posterior predicted probabilities across sampling periods (Supplementary Table 3).

### Within-Individual Consistency in Behavioral Responses

To assess within-individual consistency in behavioral responses to the aversive stimulus, behavioral responses were coded based on the discrete behaviors observed during each trial (chemical release, freezing, fleeing, escaping, bobbing, leg tapping, and autotomy). For each trial, the presence or absence of each behavior was coded as a binary variable (1 = present, 0 = absent). Within-individual behavioral variability across trials was quantified using Shannon entropy (e.g., Stamps et al. 2013), calculated in R using the diversity function in the package *vegan* (Oksanen et al. 2018).

### Behavioral Responses Modeling

Behavioral responses were analyzed using a Bayesian multivariate generalized linear mixed model. The response consisted of five behaviors: flee, escape, bobbing, chemical release, and freeze (Table 1). Each variable was modeled with a Bernoulli distribution and a logit link. Data were formatted so that each row represented a unique individual-by-sampling-period observation, with the five behaviors forming the multivariate response. Behaviors were not mutually exclusive (i.e., multiple behaviors could occur within the 5-second observation period), so the model calculated the posterior probability of performing each behavior separately. In 66% of observations (of 1070 observations of the 107 individuals), the individual performed more than one behavior. Given the complexity of accounting for the number and sequence of behaviors, and in the interest of making population-level inferences, we modeled the probability of each behavior as independent of the others.

Candidate fixed effects included sampling period, treatment food substrate (i.e., the substrate associated with the rewarding food stimulus: rock or sand), number of legs, and sensory leg condition. Unlike the substrate-use analyses, no interaction models were included for behavioral responses. In total, 12 competing models were fitted: one full model containing all biological predictors, one sampling-period-only model, three single-predictor models, three sampling-period-plus-single-predictor models, one full model without sampling period, and three leave-one-predictor-out models. Sampling period was the strongest predictor. All models excluding it had the lowest predictive support, whereas models adding food substrate, number of legs, or sensory leg condition to sampling period performed similarly to the sampling-period-only model. The full model ranked lower relative to the top-ranked model, which included sampling period and food substrate (see Results and Supplementary Table 4). We nonetheless used the full model for biological inference, to estimate the effect of each a priori predictor while accounting for the others, and we interpret the effects of the non-temporal predictors cautiously given their limited predictive contribution.

### Ethics Statement

Permits to conduct this research were provided by the Costa Rican Government (permits number: R-032-2025 and R-025-2026-OT-CONAGEBIO). This research was additionally approved by Lincoln Park Zoo’s Research Committee, the governing body that oversees all research conducted by staff of the institution. We sought approval from UIC’s Institutional Animal Care and Use Committee (IACUC), and it was waived. All animals were returned to their site of collection following the end of this study. We followed the guidelines for the use of animals in research of the Association for the Study of Animal Behavior (ASAB 2025) and the Insect Welfare Research Society (Fischer et al. 2026), as well as handling guidelines available for this taxon we had previously developed (Escalante and O’Brien 2024).

## Results

### Substrate Use During Acclimation versus Training Phases

We found no evidence that animals consistently reduced their use of the substrate associated with the aversive stimulus during training, nor that the animals increased their use of the substrate associated with food during training. Including the experimental phase by food substrate interaction did not improve predictive performance relative to the full additive model (Supplementary Table 1), providing little evidence that changes in substrate use depended on which substrate was associated with the rewarding versus aversive stimulus. Across sampling periods, individuals increasingly used other substrates rather than the substrates associated with rewarding or aversive stimuli (Fig. 2, Supplementary Table 3).

**Figure 2.**
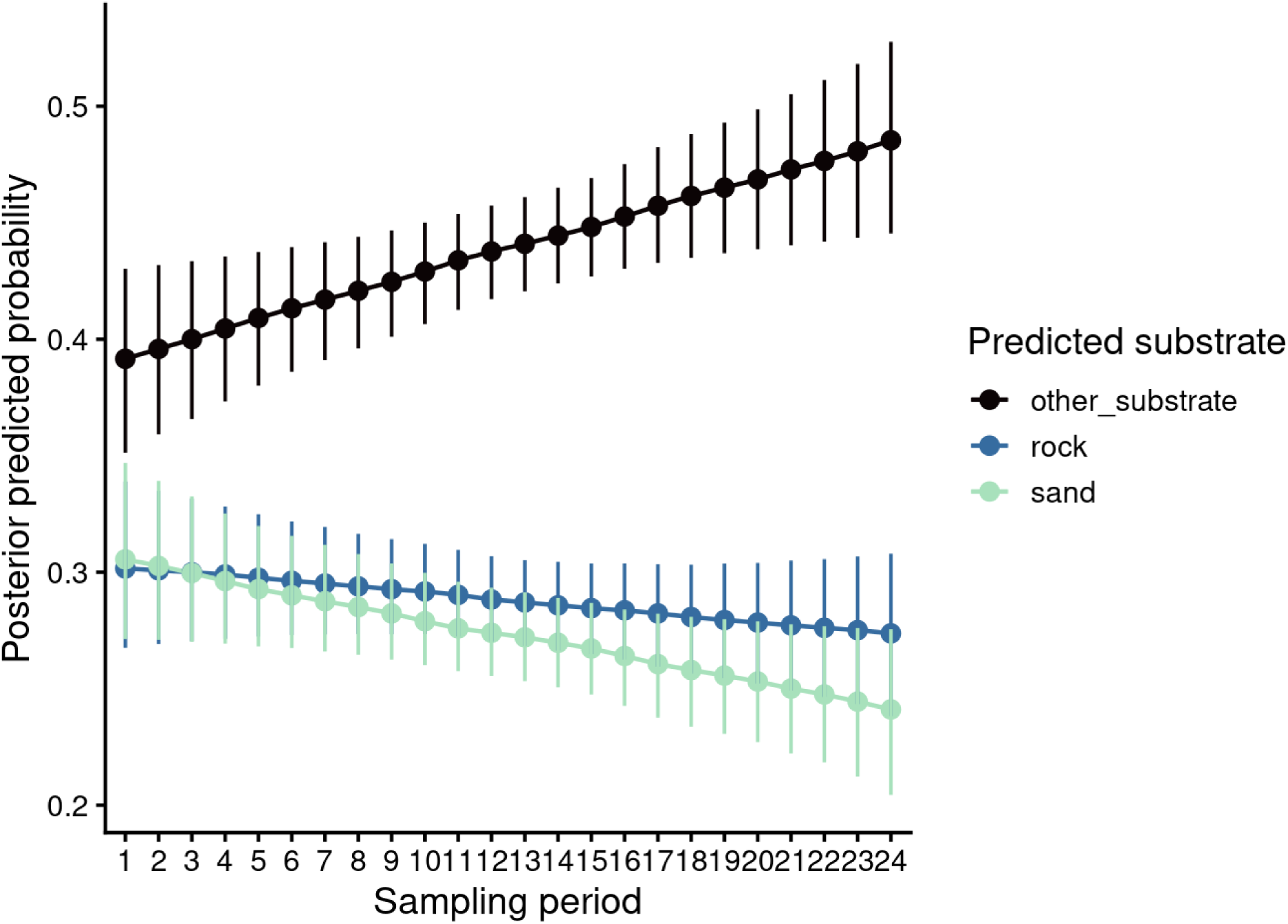
Predicted probabilities of substrate use of Opiliones across sampling periods.

### Substrate Use Between Intact versus Autotomized Individuals

Substrate use did not differ detectably between intact and autotomized animals (Fig. 3, Supplementary Table 2). Including the three-way experimental phase by substrate by leg condition interaction did not improve predictive performance (Supplementary Table 1), providing no evidence of a relationship between experimental phase, substrate use, and whether animals had eight, seven, or six legs (i.e., if they were intact or autotomized). Similarly, the presence or absence of sensory legs (both present, 1 intact/1 missing, or both missing) did not affect substrate use (Supplementary Table 2).

**Figure 3.**
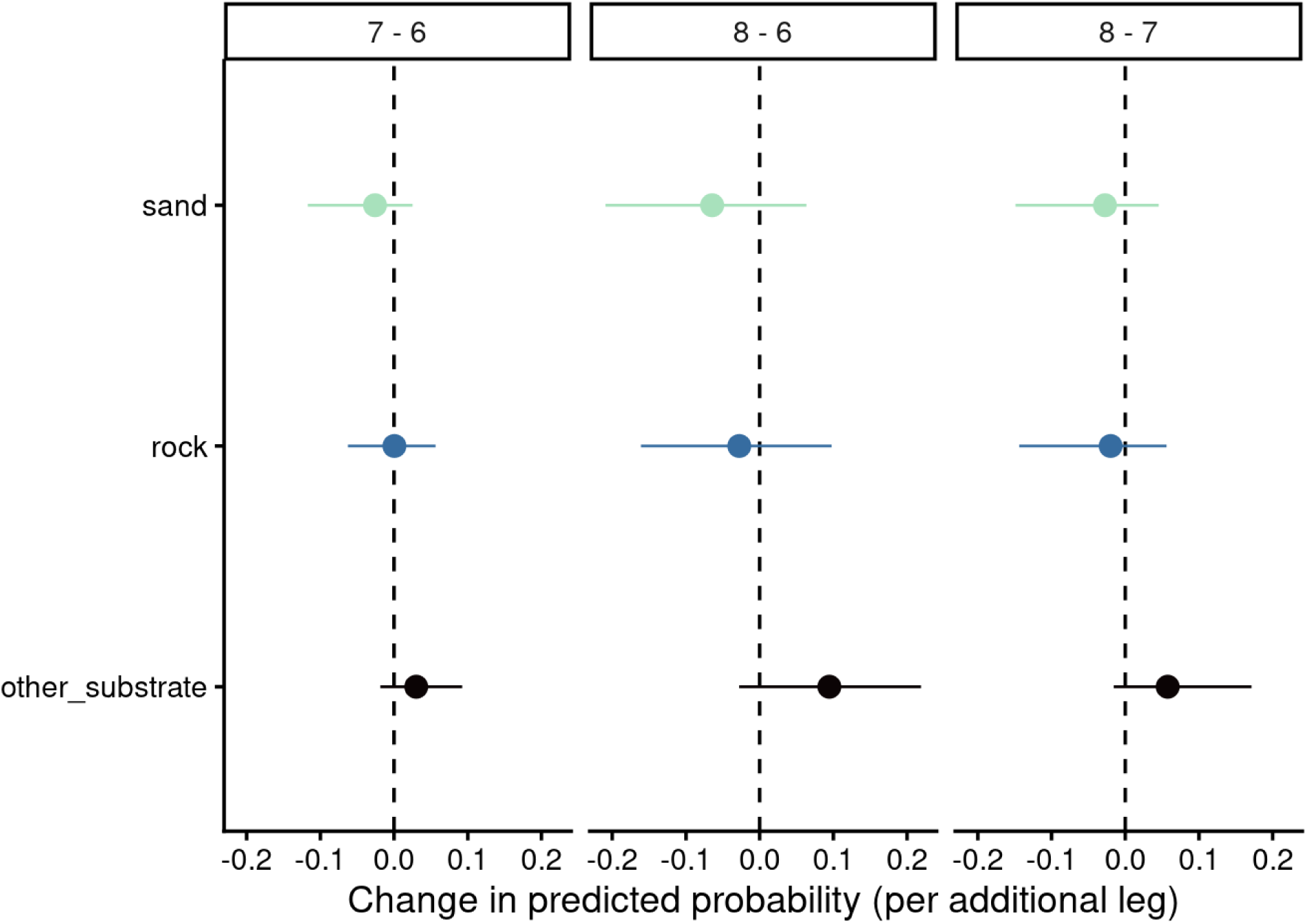
Predicted probabilities of substrate use of Opiliones based on leg condition. The left panel compares individuals with 7 legs to individuals with 6 legs, the middle panel compares individuals with 8 legs to individuals with 6 legs, and the right panel compares individuals with 8 legs to individuals with 7 legs.

### Behavioral Responses to an Aversive Stimulus

Following an aversive stimulus, flee was the most commonly observed response (42.6% of the total 1070 observations). Bobbing (17.5%), escape (14.7%), chemical release (12.6%), and freezing (12.6%) were also frequently observed. Leg tapping (0.3%) and autotomy (0.2%) were rarely observed following the aversive stimulus. Individuals generally exhibited substantial within-individual variation in their behavioral responses across repeated exposures to the aversive stimulus. Shannon entropy (*H*), where lower values indicate greater behavioral consistency and higher values indicate greater behavioral variability, had a median of 1.47 (IQR = 1.23–1.68; mean = 1.40) across individuals. Overall, these results indicate that individuals tended to vary their behavioral responses across repeated trials rather than consistently performing one behavior.

### Behavioral Responses During Acclimation versus Training Phases

Sampling period was the strongest predictor of behavioral responses, as all models excluding it had the lowest predictive support (Supplementary Table 4). Sampling period was the strongest predictor of behavioral responses, as all models excluding it had the lowest predictive support (Supplementary Table 5). Behavioral responses changed systematically over the course of the experiment, although the magnitude and direction of these changes differed among behaviors (Fig. 4, Supplementary Table 5). Flee was the most common behavioral response throughout the experiment, with a predicted probability of approximately 0.80, and exhibited little temporal variation (Fig. 4). Similarly, chemical release remained relatively stable across sampling periods, with only minor fluctuations around a predicted probability of 0.26. Freeze also showed little evidence of temporal change, maintaining a consistently low predicted probability throughout the experiment. In contrast, bobbing increased steadily across sampling periods, with the predicted probability rising from approximately 0.30 at the beginning of the experiment to about 0.42 by the final sampling period (Fig. 4). Conversely, escape declined progressively over time from approximately 0.40 to 0.24, indicating this response became less likely as the experiment progressed. Overall, temporal changes in behavioral responses were primarily driven by a gradual shift from escape toward bobbing, whereas the probabilities of flee, chemical release, and freeze remained largely unchanged.

**Figure 4.**
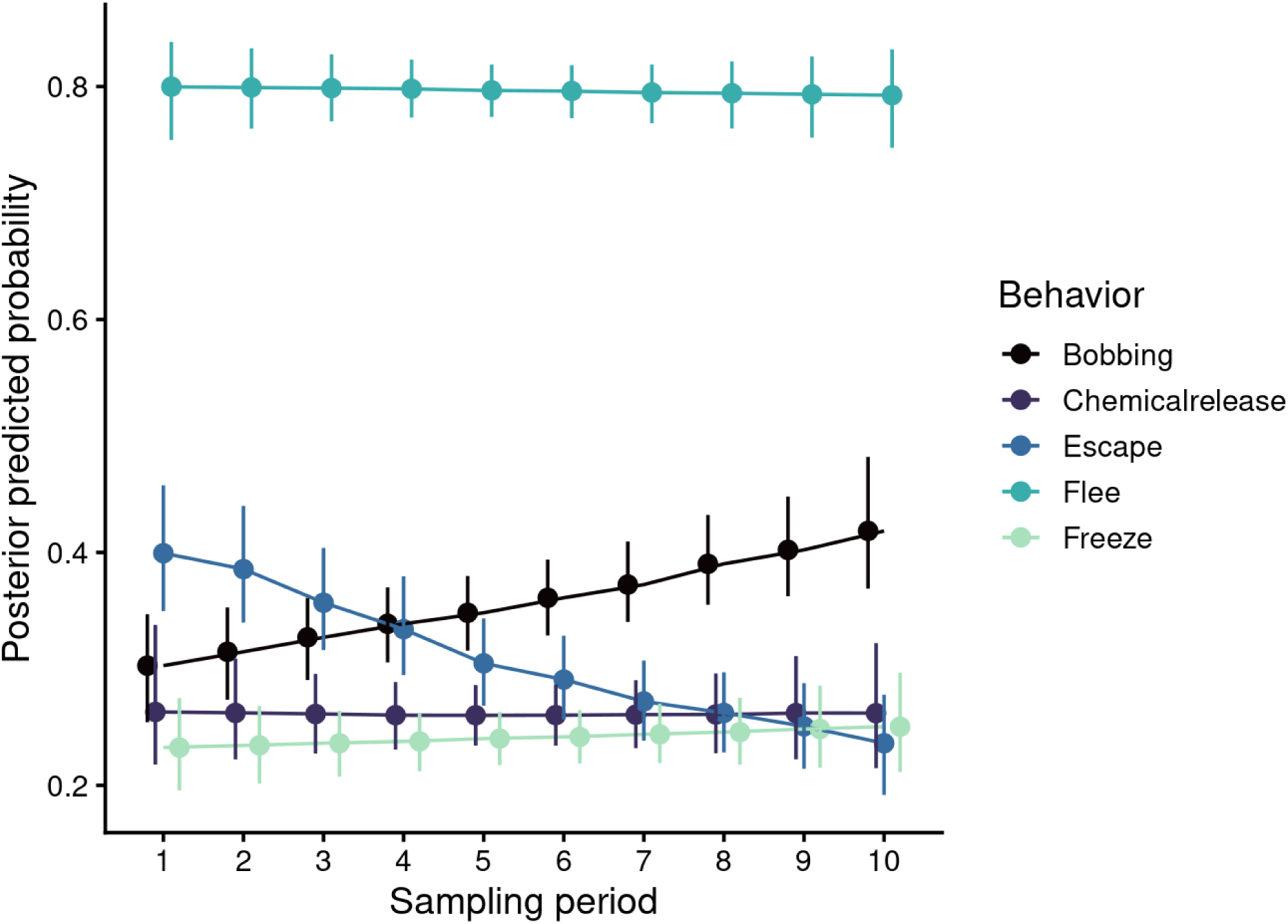
Predicted probabilities of behavioral responses of Opiliones to an aversive stimulus across sampling periods.

### Behavioral Responses Between Intact and Autotomized Individuals

Differences in behavioral responses among leg conditions were small and uncertain, and number of legs contributed little to predictive performance (Supplementary Table 4 and 6, Fig. 5). Escape showed the largest difference across leg conditions. Individuals with eight legs tended to have a higher predicted probability of escape than individuals with six legs (difference = 0.18, 95% CrI: −0.001 to 0.33) and seven legs (0.08, 95% CrI: −0.0003 to 0.25), and seven-legged individuals tended to escape more than six-legged individuals (0.08, 95% CrI: −0.0002 to 0.19), although all three credible intervals slightly overlapped zero. Flee showed little difference based on leg condition. Bobbing and freeze tended to be more likely among six-legged individuals, but these differences were small and uncertain. Chemical release varied little based on leg condition. Additionally, there was no evidence that sensory leg loss altered the behavioral repertoire of individuals (Supplementary Table 6).

**Figure 5.**
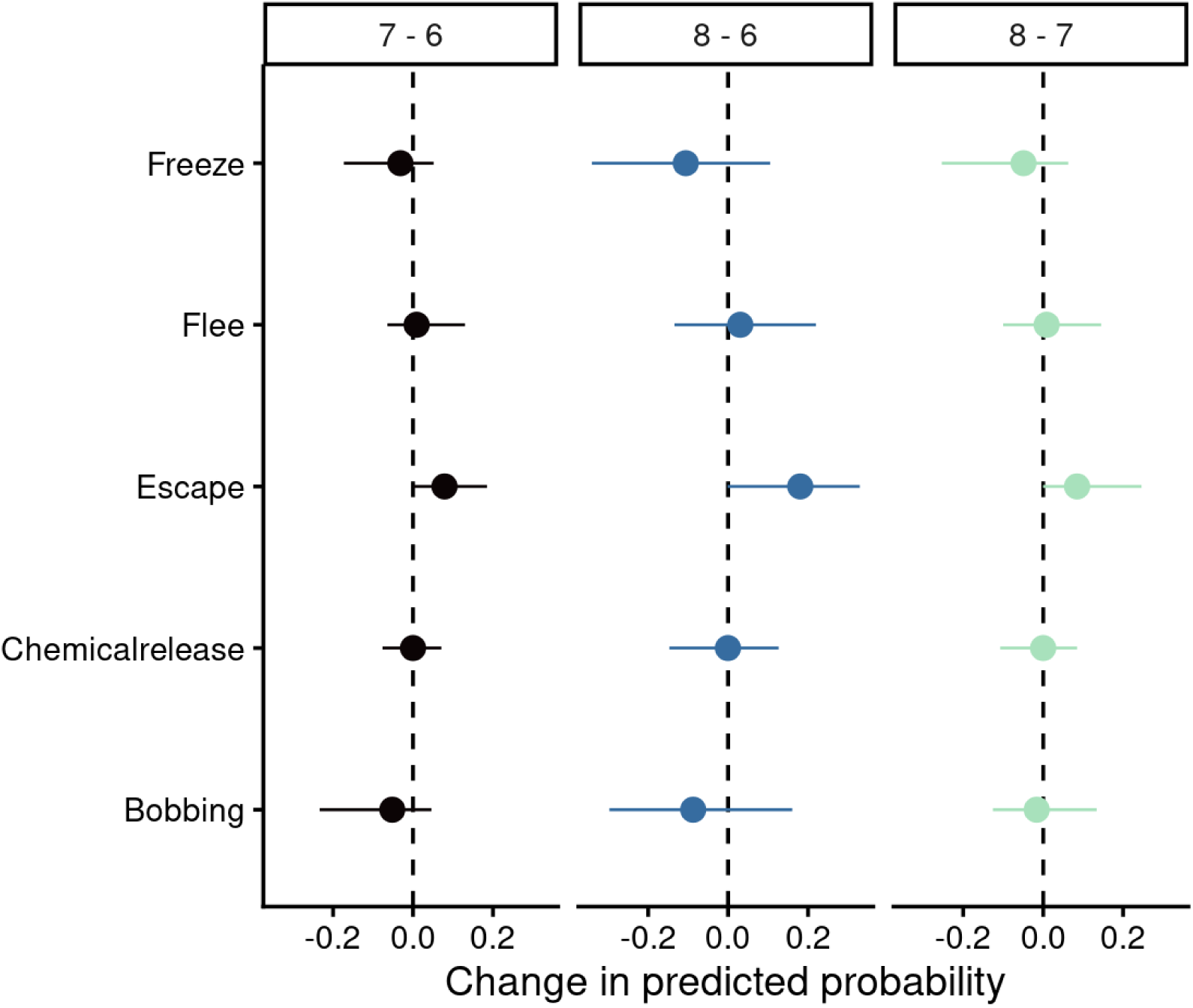
Predicted probabilities of behavioral responses of Opiliones to an aversive stimulus based on leg condition. The number of legs is denoted in the legend by color; black compares individuals with 7 legs to individuals with 6 legs, blue compares individuals with 8 legs to individuals with 6 legs, and green compares individuals with 8 legs to individuals with 7 legs.

## Discussion

### Behavioral responses to aversive stimuli

Overall, we found that all defensive behaviors observed in Opiliones of *Prionostemma* sp.1 were exhibited by both intact and autotomized individuals, although their relative frequencies tended to vary with leg condition. Eight-legged individuals were more likely to escape, whereas bobbing and freezing were more likely among individuals that had previously lost legs. These differences may reflect variation in the costs and benefits of defensive strategies according to an individual’s physical condition. For example, escaping may be more effective or less energetically costly for individuals with all legs intact (as found in Escalante et al. 2021), whereas individuals missing legs may rely more heavily on alternative defensive strategies to mitigate predation or other perceived risks. Consistent with this interpretation, Escalante et al. (2020) found that leg loss altered the frequency of locomotor gaits that might provide a defensive advantage. Furthermore, the relatively high Shannon entropy observed suggests that individuals exhibited a diversity of defensive responses across trials rather than consistently relying on a single behavioral strategy. This within-individual variation may reflect flexibility in how individuals respond to aversive experiences, potentially allowing them to employ different defensive strategies depending on their physical condition or circumstances. Together with evidence that this species avoids roosting on substrates containing chemical cues of toads, a common predator (Esquivel-Román et al., *in revision*), our findings suggest that Opiliones exhibit behavioral variation associated with both environmental threats and individual physical condition. Such context-dependent behavioral flexibility has been proposed as one line of evidence relevant to assessments of animal sentience (Birch et al. 2021; Crump et al. 2022). However, behavioral variation alone does not establish the presence of subjective experience.

Notably, autotomy was rarely observed as a response to the adverse stimulus in our study. Given the documented costs associated with autotomy (Maginnis 2006, Emberts et al. 2019, Jobson et al. 2024), leg loss may be employed as a “last resort” strategy, and/or when the immediate benefits of escaping a threat outweigh the potential longer-term consequences of losing a limb. Its infrequent occurrence may therefore reflect the relatively mild nature of the aversive stimulus used in our study, which, unlike the methods employed in previous research (DeLong et al. *in review*; Escalante et al. 2020; Escalante and Elias 2021, 2022), was not designed to induce leg release. However, the likelihood of autotomy may also depend on the nature and location of physical contact, rather than solely on the intensity of an aversive experience. Although autotomy may represent a response to particularly intense threats, further research can determine whether its occurrence reliably corresponds to more extreme negative welfare states. Given the high prevalence of leg loss in natural Opiliones populations (Escalante and O’Brien 2024), including our study species (Escalante et al. 2020), understanding how individuals adjust their defensive strategies following autotomy will inform how they navigate environmental challenges and whether particular conditions disproportionately affect individuals with different physical conditions.

### Toward validating welfare indicators

Our findings suggest that several behaviors may serve as candidate indicators of welfare in Opiliones, although their utility as indicators may depend on the duration of exposure to aversive stimuli and individual body condition. Fleeing was consistently the most common response to the aversive stimulus. As an immediate avoidance response, fleeing may be a particularly useful indicator of acute negative welfare states. In contrast, bobbing increased while escaping decreased across repeated exposures, suggesting that different behaviors may be sensitive to different aspects of aversive experiences. These contrasting trajectories could reflect changes in the relative use of defensive strategies as individuals gain experience with the stimulus, potentially arising through processes such as habituation, sensitization, or changes in arousal. In particular, increasing bobbing may be informative as a candidate indicator of responses to repeated or cumulative aversive exposure. However, further research is needed to determine whether these changes reflect sustained negative affect or other behavioral and physiological processes associated with repeated stimulation.

Notably, leg tapping (Table 1) was rarely observed following exposure to the aversive stimulus. As tapping has an exploratory or sensory function in Opiliones (Willemart et al. 2009), its relative absence under aversive conditions suggests that it may warrant investigation as an indicator of neutral or positive welfare states. Exploratory behaviors have been proposed as potential indicators of positive welfare (Crump et al. 2026; Mellor 2015), particularly when they reflect spontaneous engagement with the environment rather than responses to immediate threats or resource needs. Future studies could investigate whether tapping in *Prionostemma* increases under rewarding or otherwise favorable conditions and whether its expression reliably distinguishes between different affective states. More broadly, controlled experiments exposing individuals to stimuli of varying intensity and duration, or assessing behavioral changes following the removal of aversive conditions, could help establish the sensitivity and specificity of these candidate indicators.

Importantly, our findings highlight the need to consider individual body condition when developing and interpreting behavioral welfare indicators. Because defensive responses varied with leg condition, the same behavioral indicator may differ in its sensitivity or expression among individuals with different physical abilities. For example, reduced escape behavior among individuals missing legs may reflect physical constraints rather than a diminished response to aversive stimuli. Thus, the absence or reduced frequency of a particular behavior may reflect limitations in an individual’s ability to perform that behavior rather than differences in motivation or affective state. Consequently, welfare assessments that rely on a narrow set of behavioral indicators may underestimate responses to environmental challenges in individuals with physical disabilities. This consideration is particularly relevant for taxa in which autotomy is common, such as Opiliones (Escalante & O’Brien 2024), and those that experience substantial variation in physical condition. Validating behavioral welfare indicators across different body conditions will contribute to the development of reliable and inclusive approaches to welfare assessment.

Several limitations should be considered when interpreting our findings. Although exposure to an aversive stimulus elicited measurable defensive responses, the relationship between these behaviors and underlying affective states remains uncertain. Defensive behaviors may be mediated by nociceptive or other processes that do not necessarily require conscious perception, and their occurrence alone does not establish a negative affective experience. In the absence of independent measures of welfare, we cannot determine whether the observed responses reflect negative affect, immediate defensive reactions, or other behavioral and physiological processes. Furthermore, because individuals differed naturally in leg condition, the observed behavioral differences may reflect pre-existing individual characteristics in addition to the consequences of leg loss. Finally, responses to brief aversive stimulation under laboratory conditions may not directly correspond to responses to environmental challenges in natural settings or indicate longer-term welfare states. Although we identified potentially informative behavioral traits that can indicate the welfare of these arachnids, future research across multiple experimental contexts will establish the validity and generalizability of these candidate indicators.

### Associative learning

We found no evidence that Opiliones preferentially roosted on the substrate associated with the rewarding stimulus or avoided the substrate associated with the aversive stimulus. Instead, individuals increasingly used neutral substrates across sampling periods, suggesting that substrate selection and use may have been influenced by factors other than the experimentally established stimulus associations. These factors can include preferences for particular microhabitat characteristics or changes in roosting behavior as individuals habituated to the experimental environment. Substrate use also did not differ according to leg condition, including the number (one or two) and type (sensory versus locomotor) of missing legs. Although previous studies have demonstrated learning abilities in Opiliones (Pagoti et al. 2025; dos Santos et al. 2013), our findings did not provide evidence of associative learning under the experimental conditions.

One possible explanation is that the rock and sand substrates did not provide sufficiently distinct or ecologically relevant cues for individuals to form or express learned associations. Opiliones rely heavily on tactile and chemical information to navigate their environments (Escalante 2019, Willemart et al. 2009). While rock and sand were selected based on textural differences, these substrates may not have provided sufficiently distinctive tactile cues to the Opiliones tested. Chemical cues may therefore offer a promising alternative for investigating learning in this taxon. Additionally, the relatively mild nature of the aversive stimulus and potential variation in motivation to obtain the food reward may have limited the strength or expression of learned associations. The increasing use of neutral substrates may also reflect preferences for alternative roosting locations, potentially obscuring any learned associations with the experimental substrates. Thus, the absence of an observed substrate preference does not necessarily indicate an absence of learning, as individuals may acquire associations without expressing them through the behavioral measures assessed.

Similar findings in other understudied arachnids highlight the importance of experimental design when investigating learning. For example, Segura-Hernández et al. (2026) found that evidence of learning in Amblypygi depended on the experimental paradigm, with individuals demonstrating the ability to associate chemical cues with refuge locations despite failing to show learning in other tasks. Future studies could employ controlled choice paradigms incorporating chemical cues to investigate whether Opiliones can learn associations between environmental cues and conditions with potential welfare implications, such as barren versus enriched environments, and whether these learned associations influence decision-making across individuals with different leg conditions. Such approaches can clarify how sensory ecology and physical condition influence learning and environmental preferences, while advancing our understanding of the relationship between decision-making and welfare in Opiliones.

## Conclusion

We identified candidate behavioral indicators of welfare in the arachnid order Opiliones, a taxon for which welfare research remains limited. Fleeing, escaping, and bobbing emerged as potential indicators of negative welfare states, while exploratory behaviors such as leg tapping warrant further investigation as possible indicators of neutral or positive welfare states. The differences we found in defensive responses across repeated exposures and leg conditions suggest behavioral flexibility, highlighting how responses to aversive experiences may vary with individuals’ previous experiences and physical condition. These findings underscore the importance of accounting for individual differences in morphological traits and bodily condition when developing and interpreting behavioral welfare indicators, as indicators validated in intact individuals may not be equally informative for those that have experienced limb loss or other physical disabilities. Although we found no evidence of associative learning in the form of preferring to roost on a given substrate under the experimental conditions, future studies using alternative approaches may provide further insight into how Opiliones perceive and respond to environmental cues. Additional research is also needed to establish whether the identified behaviors reliably reflect underlying affective states and to validate their utility as welfare indicators across phenotypical and environmental contexts. Understanding how behavioral responses vary with body condition has implications beyond Opiliones, particularly for species that commonly experience autotomy, injury, or other physical changes. Ultimately, developing species-specific, empirically validated behavioral welfare indicators will be essential for advancing welfare assessment across understudied invertebrate taxa in both wild and managed environments.

## Acknowledgements

We thank the Wild Animal Initiative for their feedback on the development of this project. We also thank E. Castro, D. Brenes, G. Brenes, O. Vargas, and many other members of the staff at La Selva Research Station, and E. Castro for his assistance with the research permits. J. Ugalde provided transportation support in Costa Rica. We are very grateful to members of the University of Illinois – Chicago’s Biological Sciences Department, especially E. Beaufort, P. Ramírez, L. Lipardo, and C. Lye, for supporting the logistical and financial aspects of this project. We thank the Lincoln Park Zoo Research Committee for their support.

## Data availability statement

The data and R code supporting this study are openly available at https://github.com/maRce10/feeding-behavior-covariates-harvestmen and archived on Zenodo, https://doi.org/10.5281/zenodo.23748608). The repository contains the raw substrate-use and behavioral data, the Quarto document used to fit all models and produce the figures and supplementary tables, and a rendered version of the analysis (https://marce10.github.io/feeding-behavior-covariates-harvestmen/). A rendered version of the full analysis, including the model selection results, posterior contrasts and supplementary tables, can be viewed at https://marce10.github.io/feeding-behavior-covariates-harvestmen/. Code is released under the MIT licence and data under a CC BY 4.0 licence. Fitted model objects are not included because of their size, but they can be regenerated with the code provided.

## Author contributions

**SL O’Brien**: conceptualization (lead), funding acquisition (co-lead), data curation (lead), methodology (equal), writing – original draft (lead), writing – review and editing (equal). **M. Araya-Salas**: methodology (supporting), data curation (supporting), data analysis (lead), writing – review and editing (equal). **N. Skelly:** methodology (equal), writing – review and editing (equal). **D.M. Herrera Palencia:** methodology (equal), writing – review and editing (equal). **I. Escalante**: conceptualization (equal), methodology (equal), funding acquisition (lead), project administration (lead), supervision (lead), writing – review and editing (equal).

## Supplementary information

**Supplementary movie 1**. Examples of trials with Opiliones of *Prionostemma* sp.1 in La Selva Research Station, Sarapiquí, Costa Rica. The movie shows three individuals being tested immediately after exposure to a brief adverse stimulus (a gentle squeeze; see the description of Methods). Each clip shows and labels the behavioral response of these arachnids (as described in Table 1 above). The video can also be accessed in this link: https://drive.google.com/file/d/1ocbvUslvyT8q_AE8SLpsE_Ri1g0ubiln/view?usp=sharing

**Supplementary table 1**. Approximate leave-one-out cross-validation (PSIS-LOO) comparison of the 17 candidate models of substrate use.

**Supplementary table 2**. Posterior average contrasts in the probability of using each substrate, estimated from the full additive model.

**Supplementary table 3**. Posterior predicted probabilities of using each substrate across sampling periods, estimated from the full additive model.

**Supplementary table 4**. Approximate leave-one-out cross-validation (PSIS-LOO) comparison of the 12 candidate models of behavioral responses.

**Supplementary table 5**. Posterior predicted probabilities of each behavioral response across sampling periods, estimated from the full model.

**Supplementary table 6**. Posterior average contrasts in the probability of each behavioral response, estimated from the full model.

## Literature Cited

Acosta LE, Machado G. 2007. Diet and foraging. In: Pinto-da-Rocha R, Machado G, Giribet G, editors. Harvestmen: the biology of Opiliones. Cambridge (MA): Harvard University Press. p. 309–338.

Albalat A, Zacarias S, Coates CJ, Neil DM, Planellas SR. 2022. Welfare in farmed decapod crustaceans, with particular reference to Penaeus vannamei. Front Mar Sci 9:886024. doi:10.3389/fmars.2022.886024.

Andrade MP, Santos CMD, de Paiva MM, Medeiros SL, O’Brien CE, Lima FD, Machado JF, Leite TS. 2023. Assessing negative welfare measures for wild invertebrates: the case for octopuses. Animals 13:3021. doi:10.3390/ani13193021.

Andrews K, Birch J, Sebo J, Sims T. 2024. Background to the New York Declaration on Animal Consciousness. New York University. https://nydeclaration.com.

Animal Welfare (Sentience) Act 2022. Legislation.gov.uk, UK Public General Acts, 2022, c. 22, https://www.legislation.gov.uk/ukpga/2022/22.

Arel-Bundock V, Greifer N, Heiss A. 2024. How to interpret statistical models using marginal effects in R and Python. J Stat Softw 111:1–32. doi:10.18637/jss.v111.i09.

ASAB Ethical Committee, ABS Animal Care Committee. 2025. Guidelines for the ethical treatment of nonhuman animals in behavioural research and teaching. Anim Behav 219:123065. doi:10.1016/S0003-3472(24)00376-2.

Bacon R, Washington D, Johnson MK, Burns M. 2023. The geography of sexual conflict: a synthetic review. Am Nat 201:429–441. doi:10.1086/722797.

Barrett M, Adcock SJJ. 2023. Animal welfare science: an integral piece of sustainable insect agriculture. J Insects Food Feed 10:517–531. doi:10.1163/23524588-20230126.

Barrett M, Fischer B. 2023. Challenges in farmed insect welfare: Beyond the question of sentience. Anim Welf 32:e4. doi:10.1017/awf.2022.5.

Binding S, Farmer H, Krusin L, Cronin K. 2020. Status of animal welfare research in zoos and aquariums: where are we, where to next? J Zoo Aquar Res 8:166–174. doi:10.19227/jzar.v8i3.505.

Birch J, Burn C, Schnell A, Browning H, Crump A. 2021. Review of the evidence of sentience in cephalopod mollusks and decapod crustaceans. London: LSE Consulting, London School of Economics and Political Science. https://www.lse.ac.uk/News/News-Assets/PDFs/2021/Sentience-in-Cephalopod-Molluscs-and-Decapod-Crustaceans-Final-Report-November-2021.pdf.

Bürkner PC. 2017. brms: an R package for Bayesian multilevel models using Stan. J Stat Softw 80(1):1–28. doi:10.18637/jss.v080.i01.

Bürkner PC. 2018. Advanced Bayesian multilevel modeling with the R package brms. R J 10(1):395–411. doi:10.32614/RJ-2018-017.

Bürkner PC, Charpentier E. 2020. Modelling monotonic effects of ordinal predictors in Bayesian regression models. Br J Math Stat Psychol 73:420–451. doi:10.1111/bmsp.12195.

Brown TA, Marinko E, Burns M. 2025. The reduction of nuptial gifts in sclerosomatid Opiliones coincides with an increase in sexual conflict-like behaviour. Anim Behav 223:123150. doi:10.1016/j.anbehav.2025.123150.

Browning H, Burn C, Schnell AK, Crump A, Birch J. 2025. Animal welfare risks from commercial practices involving cephalopod mollusks and decapod crustaceans. Anim Welf 34:e24. doi:10.1017/awf.2025.25.

Calhoun AC, Tobin KB, Poorboy DM, Bowden RM, Sadd BM. 2025. To bob or not to bob: context-dependence of an antipredator response in neotropical harvestmen. Behav Ecol 36:araf004. doi:10.1093/beheco/araf004.

Cooke GM, Tonkins BM, Mather JA. 2019. Care and enrichment for captive cephalopods. In: Carere C, Mather J, editors. The welfare of invertebrate animals. Cham (Switzerland): Springer. p. 179–208. doi:10.1007/978-3-030-13947-6_8.

Crook RJ. 2021. Behavioral and neurophysiological evidence suggests affective pain experience in octopus. iScience 24:102229. doi:10.1016/j.isci.2021.102229.

Crump A, Browning H, Schnell A, Burn CC, Birch J. 2022. Sentience in decapod crustaceans: a general framework and review of the evidence. Anim Sentience 7(32):article 1. doi:10.51291/2377-7478.1691.

Crump A, Arnott G, Keeling LJ. 2026. Behavioral indicators of good welfare. In: Oldroyd B, editor. Encyclopedia of animal behavior. 3rd ed. Elsevier. p. 660–669. doi:10.1016/B978-0-443-29068-8.00228-2.

Crump A, Gibbons M, Barrett M, Birch J, Chittka L. 2023. Is it time for insect researchers to consider their subjects’ welfare? PLOS Biology 21:e3002138. doi:10.1371/journal.pbio.3002138.

de Souza Valente C. 2025. Rethinking sentience: invertebrates as worthy of moral consideration. J Agric Environ Ethics 38:3. doi:10.1007/s10806-024-09940-2.

DeLong SA, Skelly N, Freire M, Escalante I. In review. Multiple defenses, different rules: context-dependent use of chemical secretions and leg loss in an understudied arachnid. Anim Behav. Submitted 15 September 2026.

Dias BC, da Silva Souza E, Hara MR, Willemart RH. 2014. Intense leg tapping behavior by the harvestman Mischonyx cuspidatus (Gonyleptidae): an undescribed defensive behavior in Opiliones? J Arachnol 41:123–125. https://www.jstor.org/stable/24365288.

dos Santos GC, Hogan JA, Willemart RH. 2013. Associative learning in a harvestman (Arachnida, Opiliones). Behav Processes 100:64–66. doi:10.1016/j.beproc.2013.07.021.

Emberts Z, Escalante I, Bateman W. 2019. The ecology and evolution of autotomy. Biol Rev 94:1881–1896. doi:10.1111/brv.12539.

Escalante I. 2019. Chemical communication and ecology in Eupnoi harvestmen. PeerJ Preprints 7:e27778v1 doi:10.7287/peerj.preprints.27778v1.

Escalante I, Badger MA, Elias DO. 2019. Variation in movement: multiple locomotor gaits in Neotropical harvestmen. Biol J Linn Soc 127:493–507. doi:10.1093/biolinnean/blz047.

Escalante I, Badger MA, Elias DO. 2020. Rapid recovery of locomotor performance after leg loss in harvestmen. Sci Rep 10:13747. doi:10.1038/s41598-020-70557-2.

Escalante I, Elias DO. 2021. The type of leg lost affects habitat use but not survival in a non-regenerating arthropod. Ecol Evol 11:10672–10685. doi:10.1002/ece3.7879.

Escalante I, Elias DO. 2022. Robustness in action: Leg loss does not affect mating success in male harvestmen. Behav Ecol Sociobiol 76:20. doi:10.1007/s00265-022-03127-3.

Escalante I, Ellis VR, Elias DO. 2021. Leg loss decreases endurance and increases oxygen consumption during locomotion in harvestmen. J Comp Physiol A 207:257–268. doi:10.1007/s00359-020-01455-1.

Escalante I, O’Brien SL. 2024. Robustness to leg loss in Opiliones: a review and framework considerations for future research. Integr Comp Biol 64:1338–1353. doi:10.1093/icb/icae051.

Esquivel-Román A, Grullón-Peña FC, Stynoski JL, Escalante I. In revision. Tastes like danger: avoidance of a substrate with predator chemical cues in an arachnid. Behav Processes. Second revision, 1 October 2026. Associated dataset: doi:10.5061/dryad.4mw6m90r1.

Farm Animal Welfare Council. 1993. Second report on priorities for research and development in farm animal welfare. London (UK): Ministry of Agriculture, Fisheries and Food.

Fischer B, Barrett M, Adcock S, Browning H, Chittka L, Crump A, Drinkwater E, Dunlap A, Gibbons M, Perl C. 2026. Guidelines for the care and use of insects in research. Insect Welfare Research Society.

Fischer B, Gottlieb J, Schnell AK, Barrett M. 2025. Defending and refining the Birch et al. (2021) precautionary framework for animal sentience. Anim Welf 34:e28. doi:10.1017/awf.2025.27.

Fossati P, Stefanini FM, Ravasio G, Coerezza U. 2024. Welfare of invertebrates: a pilot study on a new land snail stunning technique. Sci Rep 14:8378. doi:10.1038/s41598-024-58133-4.

Fowler-Finn KD, Triana E, Miller OG. 2014. Mating in the harvestman Leiobunum vittatum (Arachnida: Opiliones): from premating struggles to solicitous tactile engagement. Behaviour 151:1663–1686.

Free D, Wolfensohn S. 2023. Assessing the welfare of captive group-housed cockroaches, Gromphadorhina oblongonota. Animals 13:3351. doi:10.3390/ani13213351.

Freire M, Skelly N, Prachand SV, Escalante I. 2026. Chemical secretion use varies among species but lacks a straightforward relationship with other defenses in Neotropical Opiliones. Ecol Evol 16:e74037. doi:10.1002/ece3.74037.

Galpayage Dona HS, Solvi C, Kowalewska A, Mäkelä K, MaBouDi H, Chittka L. 2022. Do bumble bees play? Anim Behav 194:239–251. doi:10.1016/j.anbehav.2022.08.013.

Giribet G, Sharma PP. 2015. Evolutionary biology of harvestmen (Arachnida, Opiliones). Annu Rev Entomol 60:157–175. doi:10.1146/annurev-ento-010814-021028.

Guffey C. 1999. Costs associated with leg autotomy in the harvestmen Leiobunum nigripes and Leiobunum vittatum (Arachnida: Opiliones). Can J Zool 77:824–830. doi:10.1139/z99-026.

Holmberg RG, Angerilli NPD, LaCasse LJ. 1984. Overwintering aggregations of Leiobunum paessleri in caves and mines (Arachnida, Opiliones). J Arachnol 12:195–204.

Jobson S, Hamel JF, Mercier A. 2024. Shake it off: exploring drivers and outcomes of autotomy in marine invertebrates. Biol Lett 20:20240015. doi:10.1098/rsbl.2024.0015.

Justice WSM, O’Brien MF, Szyszka O, Shotton J, Gilmour JEM, Riordan P, Wolfensohn S. 2017. Adaptation of the animal welfare assessment grid (AWAG) for monitoring animal welfare in zoological collections. Vet Rec 181:143. doi:10.1136/vr.104309.

Keller M. 2017. Feeding live invertebrate prey in zoos and aquaria: are there welfare concerns? Zoo Biol 36:316–322. doi:10.1002/zoo.21378.

Kinoshita DA, Bezerra WCA, Del Corso TM, Machado G. 2026. Going out on a limb: appendage length and function shape leg loss without evidence of autotomy modulation. Ethology 1–9. doi:10.1111/eth.70118.

Lewbart GA, Zachariah TT. 2023. Aquatic and Terrestrial Invertebrate Welfare. Animals 13:3375. doi:10.3390/ani13213375.

Machado G, Burns M. 2024. Reproductive biology of harvestmen (Arachnida: Opiliones): a review of a rapidly evolving research field. Curr Zool 70:115–135. doi:10.1093/cz/zoac102.

Maginnis TL. 2006. The costs of autotomy and regeneration in animals: a review and framework for future research, Behav Ecol 7(5):857–872. doi:10.1093/beheco/arl010.

Mellor DJ. 2015. Enhancing animal welfare by creating opportunities for positive affective engagement. N Z Vet J 63:3–8.

Mellor DJ, Beausoleil NJ, Littlewood KE, McLean AN, McGreevy PD, Jones B, Wilkins C. 2020. The 2020 five domains model: including human–animal interactions in assessments of animal welfare. Animals 10:1870. doi:10.3390/ani10101870.

Narshi TM, Free D, Justice WSM, Smith SJ, Wolfensohn S. 2022. Welfare assessment of invertebrates: adapting the animal welfare assessment grid (AWAG) for zoo decapods and cephalopods. Animals 12:1675. doi:10.3390/ani12131675.

O’Brien SL, Fischer B, Barrett M. 2026. Beyond the exhibit: addressing the welfare of feeder animals in zoological institutions. Zoo Biol. 1–7. doi:10.1002/zoo.70081.

Oksanen J, Blanchet FG, Friendly M, Kindt R, Legendre P, McGlinn D, Minchin PR, O’Hara RB, Simpson GL, Solymos P, Stevens MHH, Szoecs E, Wagner H. 2018. Vegan: community ecology package. R package version 2.5–2. https://CRAN.R-project.org/package=vegan.

Pagoti GF, Rodrigues MC, Murayama GP, Willemart RH. 2025. Aversive conditioning and memory in the harvester Mischonyx squalidus (Arachnida: Opiliones). J Arachnol 53(1):26–29. doi:10.1636/joa-s-24-005.

Pereira W, Elpino-Campos A, Del-Claro K, Machado G. 2004. Behavioral repertory of the neotropical harvestman Ilhaia cuspidata (Opiliones, Gonyleptidae). J Arachnol 32:22–30. doi:10.1636/S02-35.

Perkins K. 2021. Can aquatic invertebrates within public aquaria fit the Five Domain Welfare Model? J Appl Anim Ethics Res 3:181–204. doi:10.1163/25889567-bja10025.

Pinto-da-Rocha R, Machado G, Giribet G, editors. 2007. Harvestmen: the biology of Opiliones. Cambridge (MA): Harvard University Press.

Powell EC, Painting CJ, Hickey AJ, Machado G, Holwell GI. 2021. Diet, predators, and defensive behaviors of New Zealand harvestmen (Opiliones: Neopilionidae). J Arachnol 49:122–140. doi:10.1636/joa-s-20-002.

Proud DN, Felgenhauer BE, Townsend VR Jr, Osula DO, Gilmore WO, Napier ZL, Van Zandt PA. 2012. Diversity and habitat use of Neotropical harvestmen (Arachnida: Opiliones) in a Costa Rican rainforest. ISRN Zoology 2012:549765. doi:10.5402/2012/549765.

Raspotnig G, Hüfner A, Bodner M, Blesl J, Schaider M, Escalante I, Kunert O. 2022. Polymorphic scent gland secretions in Nelima harvestmen: “Sclerosomatid compounds” but different chemical lineages. Front Ecol Evol 10:993368. doi:10.3389/fevo.2022.993368.

Romero-González JE, Zhuo Z, Chen L, Peng C, Solvi C, Peng F. 2025. Positive affective contagion in bumble bees. Science 390:377–380. doi:10.1126/science.adr0216.

Rosenthal MF, Gertler M, Hamilton AD, Prasad S, Andrade MCB. 2017. Taxonomic bias in animal behaviour publications. Anim Behav 127:83–89. doi:10.1016/j.anbehav.2017.02.017.

Rowe E, Robles López KY, Robinson KM, Baudier KM, Barrett M. 2024. Farmed cricket (Acheta domesticus, Gryllus assimilis, and Gryllodes sigillatus; Orthoptera) welfare considerations: recommendations for improving global practice. J Insects Food Feed 10:1253–1311. doi:10.1163/23524588-00001087.

Sasson DA, Johnson TD, Scott ER, Fowler-Finn KD. 2020. Short-term water deprivation has widespread effects on mating behaviour in a harvestman. Anim Behav 165:97–106. doi:10.1016/j.anbehav.2020.04.026.

Schoffstall B, Morales N, Montague J. 2013. Note on photographing embryos and hatchlings of Leiobunum sp. Harvestmen (Arachnida: Opiliones: Sclerosomatidae: Leiobuninae). Fla Entomol 96:679–681. doi:10.1653/024.096.0246.

Segura-Hernández L, Lehmann KD, Perez J, Wiegmann DD, Bingman VP, Hebets EA. 2026. Olfactory learning in two Amblypygi species Paraphrynus laevifrons and Phrynus pseudoparvulus. Anim Cogn 29:44. doi:10.1007/s10071-026-02065-1.

Stamps JA, Saltz JB, Krishnan VV. 2013. Genotypic differences in behavioural entropy: unpredictable genotypes are composed of unpredictable individuals. Anim Behav 86:641–649. doi:10.1016/j.anbehav.2013.07.012.

Stan Development Team. 2021. Stan modeling language users guide and reference manual. Version 2.26. https://mc-stan.org/docs/2_26/stan-users-guide/.

Vehtari A, Gabry J, Magnusson M, Yao Y, Bürkner PC, Paananen T, Gelman A. 2025. loo: efficient leave-one-out cross-validation and WAIC for Bayesian models. R package version 2.10.1. https://mc-stan.org/loo/.

Vehtari A, Gelman A, Gabry J. 2017. Practical Bayesian model evaluation using leave-one-out cross-validation and WAIC. Stat Comput 27:1413–1432. doi:10.1007/s11222-016-9696-4.

Vehtari A, Simpson D, Gelman A, Yao Y, Gabry J. 2024. Pareto smoothed importance sampling. J Mach Learn Res 25:1–58. https://www.jmlr.org/papers/v25/19-556.html.

Villaseñor-Amador D, Escalante I. 2025. Grouped and vibrating: the influence of group size, triggering stimuli, and leg loss in bobbing in an arachnid. Ethology 131:e13551. doi:10.1111/eth.13551.

Villaseñor-Amador D, Escalante I. 2026. When missing legs, Prionostemma Opiliones roost more frequently in the aggregation’s centers. J Arachnol 54:1–5. doi:10.1636/JoA-S-24-043.

Wahltinez SJ, Stacy NI, Hadfield CA, Harms CA, Lewbart GA, Newton AL, Nunamaker EA. 2022. Perspective: Opportunities for advancing aquatic invertebrate welfare. Front Vet Sci 9:973376. doi:10.3389/fvets.2022.973376.

Willemart RH, Farine JP, Peretti AV, Gnaspini P. 2006. Behavioral roles of the sexually dimorphic structures in the male harvestman, Phalangium opilio (Opiliones, Phalangiidae). Can J Zool 84:1763–1774. doi:10.1139/z06-173.

Willemart RH, Farine JP, Gnaspini P. 2009. Sensory biology of Phalangida harvestmen (Arachnida, Opiliones): a review, with new morphological data on 18 species. Acta Zool 90:209–227. doi:10.1111/j.1463-6395.2008.00341.x.

Wilson-Sanders SE. 2011. Invertebrate Models for Biomedical Research, Testing, and Education. ILAR J 52:126–152. doi:10.1093/ilar.52.2.126.

Ximenes NG, Willemart RH. 2024. Staying alive: how harvesters avoid predation. J Arachnol 52:127–138. doi:10.1636/JoA-S-22-054.

