## Supplemental Tables 1-6 for "Toward welfare indicators: leg condition and aversive stimuli modulate defensive behavior in Opilones"

**Supplementary Table 1.** Approximate leave-one-out cross-validation (PSIS-LOO) comparison of the 17 candidate Bayesian categorical mixed models of substrate use (rock, sand or other substrate) in Opiliones. Models are ranked by expected log pointwise predictive density ($elpd_{loo}$).

| Rank | Model | Fixed effects | Δelpd | SE Δelpd | $elpd_{loo}$ (SE) | $p_{loo}$ | LOOIC | Stacking weight |
|---|---|---|---|---|---|---|---|---|
| 1 | drop_sensory_leg_missing | sampling period (mo) + field season + food substrate + number of legs (mo) + phase | 0.00 | 0.00 | -2511.95 (21.59) | 132.3 | 5023.9 | 0.098 |
| 2 | full | sampling period (mo) + field season + food substrate + number of legs (mo) + sensory leg missing + phase | -0.65 | 0.70 | -2512.60 (21.67) | 133.5 | 5025.2 | 0.000 |
| 3 | drop_number_of_legs | sampling period (mo) + field season + food substrate + sensory leg missing + phase | -0.81 | 1.11 | -2512.76 (21.56) | 132.7 | 5025.5 | 0.000 |
| 4 | full_interaction | sampling period (mo) + field season + food substrate × phase + number of legs (mo) + sensory leg missing | -1.75 | 1.95 | -2513.70 (21.77) | 135.8 | 5027.4 | 0.000 |
| 5 | single_sampling_period_food_substrate | sampling period (mo) + field season + food substrate | -2.28 | 2.96 | -2514.23 (21.29) | 129.6 | 5028.5 | 0.200 |
| 6 | drop_sampling_period | field season + food substrate + number of legs (mo) + sensory leg missing + phase | -2.97 | 2.90 | -2514.92 (21.45) | 131.2 | 5029.8 | 0.269 |
| 7 | drop_phase | sampling period (mo) + field season + food substrate + number of legs (mo) + sensory leg missing | -3.30 | 2.86 | -2515.24 (21.45) | 132.1 | 5030.5 | 0.000 |
| 8 | full_three_way_interaction | sampling period (mo) + field season + food substrate × phase × leg condition + number of legs (mo) + sensory leg missing | -3.49 | 3.58 | -2515.44 (22.12) | 138.2 | 5030.9 | 0.146 |
| 9 | single_sampling_period_phase | sampling period (mo) + field season + phase | -7.36 | 4.07 | -2519.31 (21.25) | 148.3 | 5038.6 | 0.000 |
| 10 | drop_food_substrate | sampling period (mo) + field season + number of legs (mo) + sensory leg missing + phase | -8.03 | 4.04 | -2519.97 (21.41) | 150.2 | 5039.9 | 0.000 |
| 11 | single_food_substrate | field season + food substrate | -8.76 | 5.09 | -2520.71 (20.80) | 126.3 | 5041.4 | 0.001 |
| 12 | sampling_period_only | sampling period (mo) + field season | -9.08 | 5.01 | -2521.02 (21.09) | 146.4 | 5042.0 | 0.239 |
| 13 | single_phase | field season + phase | -9.29 | 4.87 | -2521.24 (20.99) | 145.5 | 5042.5 | 0.047 |
| 14 | single_sampling_period_number_of_legs | sampling period (mo) + field season + number of legs (mo) | -9.44 | 4.92 | -2521.39 (21.16) | 147.3 | 5042.8 | 0.000 |
| 15 | single_sampling_period_sensory_leg_missing | sampling period (mo) + field season + sensory leg missing | -9.92 | 5.05 | -2521.87 (21.16) | 147.9 | 5043.7 | 0.000 |
| 16 | single_number_of_legs | field season + number of legs (mo) | -16.84 | 6.40 | -2528.79 (20.60) | 144.4 | 5057.6 | 0.000 |
| 17 | single_sensory_leg_missing | field season + sensory leg missing | -17.14 | 6.48 | -2529.09 (20.60) | 144.6 | 5058.2 | 0.000 |

*Note.*
Δelpd: difference in $elpd_{loo}$ relative to the top-ranked model; SE Δelpd: standard error of that difference; $p_{loo}$: effective number of parameters; LOOIC = $-2\times$ $elpd_{loo}$. Stacking weights were computed with loo::loo_model_weights(method = "stacking"). All models include a random intercept for individual and were fitted with brms (categorical family; 4 chains, 4,000 iterations each). (mo) denotes monotonic effects. N = 2,521 observations. All Pareto $k$ values were $< 0.7$ (maximum = 0.46).

**Supplementary Table 2.** Posterior average contrasts in the probability of using each substrate from the full additive categorical mixed model of substrate use, averaged over the observed values of all other predictors.

| Contrast | Substrate | Estimate [95% CrI] | |
|---|---|---|---|
| **Food substrate** | | | |
| sand − rock | Other substrate | -0.014 [-0.069, 0.040] | |
| sand − rock | Rock | 0.210 [0.156, 0.265] | * |
| sand − rock | Sand | -0.196 [-0.254, -0.138] | * |
| **Experimental phase** | | | |
| training − acclimation | Other substrate | 0.053 [0.012, 0.096] | * |
| training − acclimation | Rock | -0.009 [-0.047, 0.027] | |
| training − acclimation | Sand | -0.044 [-0.081, -0.007] | * |
| **Number of legs** | | | |
| 7 legs − 6 legs | Other substrate | 0.030 [-0.019, 0.093] | |
| 8 legs − 6 legs | Other substrate | 0.095 [-0.028, 0.219] | |
| 8 legs − 7 legs | Other substrate | 0.058 [-0.016, 0.171] | |
| 7 legs − 6 legs | Rock | 0.000 [-0.063, 0.056] | |
| 8 legs − 6 legs | Rock | -0.028 [-0.161, 0.098] | |
| 8 legs − 7 legs | Rock | -0.020 [-0.144, 0.056] | |
| 7 legs − 6 legs | Sand | -0.026 [-0.117, 0.025] | |
| 8 legs − 6 legs | Sand | -0.064 [-0.209, 0.064] | |
| 8 legs − 7 legs | Sand | -0.027 [-0.149, 0.045] | |
| **Sensory leg condition** | | | |
| sensory leg(s) missing − both present | Other substrate | 0.019 [-0.066, 0.104] | |
| sensory leg(s) missing − both present | Rock | -0.046 [-0.124, 0.040] | |
| sensory leg(s) missing − both present | Sand | 0.025 [-0.061, 0.118] | |

*Note.*
Estimates are posterior means of the difference in predicted probability (e.g., 0.10 = 10 percentage points) with 95% credible intervals (CrI). * 95% CrI excludes zero. Food substrate refers to the substrate on which food (rewarding stimulus) was presented.

**Supplementary Table 3.** Posterior predicted probabilities [95% CrI] of using each substrate across sampling periods (modeled as a monotonic effect) from the full additive categorical mixed model of substrate use, averaged over the observed values of all other predictors.

| Sampling period | Other substrate | Rock | Sand |
|---|---|---|---|
| 1 | 0.392 [0.351, 0.430] | 0.302 [0.268, 0.339] | 0.306 [0.271, 0.347] |
| 2 | 0.396 [0.359, 0.432] | 0.301 [0.269, 0.335] | 0.303 [0.271, 0.339] |
| 3 | 0.400 [0.366, 0.433] | 0.300 [0.270, 0.332] | 0.300 [0.270, 0.332] |
| 4 | 0.405 [0.373, 0.435] | 0.299 [0.271, 0.328] | 0.296 [0.269, 0.325] |
| 5 | 0.409 [0.380, 0.437] | 0.298 [0.272, 0.325] | 0.293 [0.268, 0.320] |
| 6 | 0.413 [0.386, 0.440] | 0.296 [0.273, 0.322] | 0.290 [0.267, 0.316] |
| 7 | 0.417 [0.391, 0.442] | 0.295 [0.273, 0.319] | 0.288 [0.266, 0.312] |
| 8 | 0.421 [0.396, 0.444] | 0.294 [0.273, 0.316] | 0.285 [0.265, 0.308] |
| 9 | 0.425 [0.401, 0.447] | 0.293 [0.273, 0.314] | 0.282 [0.263, 0.304] |
| 10 | 0.429 [0.406, 0.450] | 0.292 [0.273, 0.312] | 0.279 [0.260, 0.300] |
| 11 | 0.434 [0.413, 0.454] | 0.290 [0.272, 0.310] | 0.276 [0.257, 0.296] |
| 12 | 0.438 [0.417, 0.457] | 0.288 [0.270, 0.307] | 0.274 [0.256, 0.293] |
| 13 | 0.441 [0.421, 0.461] | 0.287 [0.268, 0.305] | 0.272 [0.253, 0.291] |
| 14 | 0.444 [0.424, 0.465] | 0.286 [0.266, 0.304] | 0.270 [0.251, 0.289] |
| 15 | 0.448 [0.427, 0.469] | 0.285 [0.265, 0.304] | 0.267 [0.248, 0.287] |
| 16 | 0.453 [0.430, 0.475] | 0.284 [0.263, 0.304] | 0.264 [0.243, 0.284] |
| 17 | 0.457 [0.433, 0.482] | 0.282 [0.259, 0.303] | 0.261 [0.238, 0.282] |
| 18 | 0.461 [0.435, 0.488] | 0.281 [0.257, 0.303] | 0.258 [0.234, 0.281] |
| 19 | 0.465 [0.437, 0.493] | 0.279 [0.255, 0.304] | 0.256 [0.231, 0.280] |
| 20 | 0.469 [0.439, 0.499] | 0.278 [0.252, 0.304] | 0.253 [0.227, 0.279] |
| 21 | 0.473 [0.440, 0.505] | 0.277 [0.249, 0.305] | 0.250 [0.222, 0.278] |
| 22 | 0.476 [0.442, 0.511] | 0.276 [0.246, 0.306] | 0.248 [0.218, 0.277] |
| 23 | 0.481 [0.444, 0.518] | 0.275 [0.243, 0.307] | 0.244 [0.212, 0.276] |
| 24 | 0.485 [0.445, 0.528] | 0.274 [0.238, 0.308] | 0.241 [0.204, 0.276] |

**Supplementary Table 4.** Approximate leave-one-out cross-validation (PSIS-LOO) comparison of the 12 candidate Bayesian multivariate Bernoulli mixed models of behavioral responses (flee, escape, bobbing, chemical release and freeze) to an aversive stimulus in Opiliones. Models are ranked by expected log pointwise predictive density ($elpd_{loo}$).

| Rank | Model | Fixed effects | Δelpd | SE Δelpd | $elpd_{loo}$ (SE) | $p_{loo}$ | LOOIC | Stacking weight |
|---|---|---|---|---|---|---|---|---|
| 1 | single__sampling__period__Treatment__food__substrate | sampling period (mo) + field season + food substrate | 0.00 | 0.00 | -2707.30 (44.58) | 337.8 | 5414.6 | 0.270 |
| 2 | sampling__period__only | sampling period (mo) + field season | -0.35 | 1.55 | -2707.65 (44.45) | 336.7 | 5415.3 | 0.000 |
| 3 | single__sampling__period__Number__of__legs | sampling period (mo) + field season + number of legs (mo) | -0.38 | 2.28 | -2707.68 (44.72) | 338.8 | 5415.4 | 0.452 |
| 4 | drop__Sensory__leg__missing | sampling period (mo) + field season + food substrate + number of legs (mo) | -1.03 | 1.80 | -2708.34 (44.85) | 340.2 | 5416.7 | 0.072 |
| 5 | drop__Number__of__legs | sampling period (mo) + field season + food substrate + sensory leg missing | -1.89 | 1.55 | -2709.19 (44.82) | 341.4 | 5418.4 | 0.005 |
| 6 | single__sampling__period__Sensory__leg__missing | sampling period (mo) + field season + sensory leg missing | -2.02 | 2.11 | -2709.32 (44.70) | 340.0 | 5418.6 | 0.000 |
| 7 | drop__Treatment__food__substrate | sampling period (mo) + field season + number of legs (mo) + sensory leg missing | -3.10 | 2.57 | -2710.40 (44.97) | 343.4 | 5420.8 | 0.001 |
| 8 | full | sampling period (mo) + field season + food substrate + number of legs (mo) + sensory leg missing | -3.84 | 2.02 | -2711.14 (45.05) | 344.7 | 5422.3 | 0.000 |
| 9 | single__Treatment__food__substrate | field season + food substrate | -9.38 | 5.69 | -2716.69 (44.13) | 328.2 | 5433.4 | 0.185 |
| 10 | single__Sensory__leg__missing | field season + sensory leg missing | -10.71 | 6.04 | -2718.01 (44.26) | 329.7 | 5436.0 | 0.014 |
| 11 | single__Number__of__legs | field season + number of legs (mo) | -10.72 | 6.03 | -2718.03 (44.31) | 330.4 | 5436.1 | 0.000 |
| 12 | drop__sampling__period | field season + food substrate + number of legs (mo) + sensory leg missing | -13.06 | 6.01 | -2720.37 (44.65) | 335.4 | 5440.7 | 0.000 |

*Note.*
Δelpd: difference in $elpd_{loo}$ relative to the top-ranked model; SE Δelpd: standard error of that difference; $p_{loo}$: effective number of parameters; LOOIC $= -2\times$ $elpd_{loo}$. Stacking weights were computed with loo::loo__model__weights(method = "stacking"). All models include a random intercept for individual and were fitted with brms (Bernoulli family with logit link for each of the five behaviors, residual correlations not modeled; predictors were identical across behaviors; 4 chains, 4,000 iterations each). (mo) denotes monotonic effects. N = 1,070 observations. All Pareto $k$ values were $< 0.7$ (maximum = 0.82).

**Supplementary Table 5.** Posterior predicted probabilities [95% CrI] of each behavioral response to the aversive stimulus across sampling periods (modeled as a monotonic effect) from the full multivariate Bernoulli mixed model of behavioral responses, averaged over the observed values of all other predictors.

| Sampling period | Bobbing | Chemical release | Escape | Flee | Freeze |
|---|---|---|---|---|---|
| 1 | 0.303 [0.254, 0.347] | 0.263 [0.218, 0.338] | 0.399 [0.350, 0.458] | 0.800 [0.754, 0.838] | 0.232 [0.196, 0.275] |
| 2 | 0.315 [0.274, 0.353] | 0.262 [0.222, 0.309] | 0.386 [0.340, 0.440] | 0.799 [0.764, 0.833] | 0.234 [0.201, 0.268] |
| 3 | 0.327 [0.290, 0.361] | 0.261 [0.227, 0.296] | 0.357 [0.316, 0.404] | 0.799 [0.770, 0.828] | 0.236 [0.208, 0.264] |
| 4 | 0.339 [0.306, 0.370] | 0.260 [0.231, 0.289] | 0.334 [0.295, 0.380] | 0.798 [0.773, 0.823] | 0.238 [0.212, 0.262] |
| 5 | 0.348 [0.316, 0.380] | 0.260 [0.234, 0.286] | 0.305 [0.268, 0.343] | 0.797 [0.774, 0.819] | 0.240 [0.217, 0.263] |
| 6 | 0.361 [0.329, 0.394] | 0.260 [0.234, 0.287] | 0.291 [0.256, 0.329] | 0.796 [0.773, 0.818] | 0.242 [0.219, 0.265] |
| 7 | 0.372 [0.340, 0.409] | 0.261 [0.232, 0.290] | 0.272 [0.238, 0.307] | 0.795 [0.768, 0.819] | 0.244 [0.219, 0.270] |
| 8 | 0.390 [0.355, 0.432] | 0.261 [0.228, 0.296] | 0.262 [0.228, 0.297] | 0.794 [0.764, 0.821] | 0.246 [0.218, 0.275] |
| 9 | 0.402 [0.362, 0.448] | 0.262 [0.222, 0.311] | 0.251 [0.214, 0.288] | 0.793 [0.756, 0.826] | 0.248 [0.215, 0.286] |
| 10 | 0.418 [0.369, 0.482] | 0.262 [0.215, 0.322] | 0.236 [0.192, 0.278] | 0.793 [0.747, 0.832] | 0.250 [0.211, 0.297] |

**Supplementary Table 6.** Posterior average contrasts in the probability of each behavioral response to the aversive stimulus from the full multivariate Bernoulli mixed model of behavioral responses, averaged over the observed values of all other predictors.

| Contrast | Behavior | Estimate [95% CrI] | |
|---|---|---|---|
| **Food substrate** | | | |
| sand − rock | Bobbing | -0.071 [-0.168, 0.028] | |
| sand − rock | Chemical release | -0.028 [-0.097, 0.042] | |
| sand − rock | Escape | -0.111 [-0.205, -0.017] | * |
| sand − rock | Flee | -0.008 [-0.089, 0.076] | |
| sand − rock | Freeze | -0.043 [-0.144, 0.060] | |
| **Number of legs** | | | |
| 7 legs − 6 legs | Bobbing | -0.052 [-0.234, 0.046] | |
| 8 legs − 6 legs | Bobbing | -0.087 [-0.297, 0.161] | |
| 8 legs − 7 legs | Bobbing | -0.016 [-0.126, 0.134] | |
| 7 legs − 6 legs | Chemical release | 0.000 [-0.076, 0.071] | |
| 8 legs − 6 legs | Chemical release | 0.000 [-0.147, 0.126] | |
| 8 legs − 7 legs | Chemical release | 0.000 [-0.107, 0.085] | |
| 7 legs − 6 legs | Escape | 0.079 [0.000, 0.185] | |
| 8 legs − 6 legs | Escape | 0.180 [-0.001, 0.330] | |
| 8 legs − 7 legs | Escape | 0.085 [0.000, 0.246] | |
| 7 legs − 6 legs | Flee | 0.009 [-0.064, 0.130] | |
| 8 legs − 6 legs | Flee | 0.030 [-0.134, 0.220] | |
| 8 legs − 7 legs | Flee | 0.009 [-0.100, 0.145] | |
| 7 legs − 6 legs | Freeze | -0.032 [-0.173, 0.052] | |
| 8 legs − 6 legs | Freeze | -0.106 [-0.341, 0.105] | |
| 8 legs − 7 legs | Freeze | -0.049 [-0.254, 0.063] | |
| **Sensory leg condition** | | | |
| sensory leg(s) missing − both present | Bobbing | -0.028 [-0.164, 0.122] | |
| sensory leg(s) missing − both present | Chemical release | -0.023 [-0.113, 0.076] | |
| sensory leg(s) missing − both present | Escape | 0.015 [-0.118, 0.168] | |
| sensory leg(s) missing − both present | Flee | -0.093 [-0.241, 0.043] | |
| sensory leg(s) missing − both present | Freeze | 0.019 [-0.119, 0.188] | |

*Note.*
Estimates are posterior means of the difference in predicted probability (e.g., 0.10 = 10 percentage points) with 95% credible intervals (CrI). * 95% CrI excludes zero. Food substrate refers to the substrate on which food (rewarding stimulus) was presented.
